# Epidermal GIGANTEA adjusts the response to shade at dusk by directly impinging on PHYTOCHROME INTERACTING FACTOR 7 function

**DOI:** 10.1101/2023.03.21.533699

**Authors:** Carlos Martínez-Vasallo, Benjamin Cole, Javier Gallego-Bartolomé, Joanne Chory, Steve A. Kay, Maria A. Nohales

**Affiliations:** Instituto de Biología Molecular y Celular de Plantas, Consejo Superior de Investigaciones Científicas - Universidad Politécnica de Valencia, 46022 Valencia, Spain; Keck School of Medicine, University of Southern California, Los Angeles, CA 90089, USA; Plant Biology Laboratory, Salk Institute for Biological Studies, 10010 North Torrey Pines Road, La Jolla, CA 92037, USA; Howard Hughes Medical Institute, Salk Institute for Biological Studies, La Jolla, CA 92037, USA

**Keywords:** Shade avoidance, circadian gating, GIGANTEA, PIF7, tissue specificity

## Abstract

For plants adapted to bright light, a decrease in the amount of light received can be detrimental to their growth and survival. Consequently, in response to shade from surrounding vegetation, they initiate a suite of molecular and morphological changes known as the shade avoidance response (SAR) through which stems and petioles elongate in search for light. Under sunlight-night cycles, the plant’s responsiveness to shade varies across the day, being maximal at dusk time. While a role for the circadian clock in this regulation has long been proposed, mechanistic understanding of how it is achieved is incomplete. Here we show that the clock component GIGANTEA (GI) directly interacts with the transcriptional regulator PHYTOCHROME INTERACTING FACTOR 7 (PIF7), a key player in the response to shade. GI represses PIF7 transcriptional activity and the expression of its target genes in response to shade, thereby fine-tuning the magnitude of the response to limiting light conditions. We find that, under light/dark cycles, this function of GI is required to adequately modulate the gating of the response to shade at dusk. Importantly, we also show that GI expression in epidermal cells is sufficient for proper SAR regulation.

**SIGNIFICANCE:** Plants have a remarkable capacity to adapt to and cope with changes in environmental conditions. Because of the importance of light to their survival, plants have evolved sophisticated mechanisms to optimize responses to light. An outstanding adaptive response in terms of plant plasticity in dynamic light environments is the shade avoidance response which sun-loving plants deploy to escape canopy and grow towards the light. This response is the result of a complex signaling network in which cues from different pathways are integrated, including light, hormone, and circadian signaling. Within this framework, our study provides a mechanistic model of how the circadian clock contributes to this complex response by temporalizing the sensitivity to shade signals towards the end of the light period. In light of evolution and local adaptation, this work gives insights into a mechanism through which plants may have optimized resource allocation in fluctuating environments.

## INTRODUCTION

Light is a key environmental cue and resource for plants, as they use it to interpret their surroundings but also rely on it to perform photosynthesis and fix carbon, which is essential for plant growth and development. In both natural and agricultural settings, the light environment is highly dynamic and plants are constantly monitoring light quantity and quality to adapt to it accordingly. Because of the importance of light to their survival, plants have evolved exquisite mechanisms to maximize exploitation of this resource and to cope with unfavorable conditions (such as limiting or high-intensity light). For plants adapted to open environments, changes in light quality caused by neighboring vegetation are interpreted as a threat entailing competition for light and trigger an adaptive response to escape canopy known as the shade avoidance response (SAR) (1). Phenotypically, the SAR comprises a series of morphological changes which include stem and petiole elongation, leaf hyponasty and early flowering, among others (1-3).

At the molecular level, the proximity of other plants is sensed as a change in the ratio of red to far-red (R:FR) light, which is caused by an enrichment in the FR wavelengths of the light spectrum that are reflected and transmitted through the leaves of the surrounding vegetation (4). This change in light quality is perceived by the phytochrome family of photoreceptors, especially phytochrome B (phyB) (4-6), which then transduce the signal to transcriptional networks through the regulation of the activity of the basic helix-loop-helix (bHLH) transcription factors PHYTOCHROME INTERACTING FACTORS (PIFs) (4). Under bright light, where the R:FR ratio is high, phyB is in its active far-red absorbing form (Pfr) which localizes in the nucleus where it physically interacts with PIFs and promotes their phosphorylation and subsequent degradation. In the shade, the enrichment in FR light (low R:FR ratio) promotes the photoconversion of Pfr to its inactive red-absorbing form (Pr) which is translocated to the cytoplasm thereby allowing PIF accumulation and activity (4). This then enables the induction of the expression of auxin biosynthesis enzymes and cell elongation genes, which promote and support shade-induced growth (7, 8). Several PIFs have been implicated in the response to shade, including PIF4 and 5 (9) and PIF7 (10, 11), which seems to play a dominant role in this pathway.

In the field, plants must adapt a sessile lifestyle under fluctuating environments and are presented with a variety of challenges on a daily basis, many of which arise from the existence of day/night cycles. These cycles generate, for example, large but predictable fluctuations in important ambient variables including light intensity. In this context, organisms have evolved circadian clocks as endogenous time tracking mechanisms that enable them to anticipate these changes and to organize their physiology accordingly, precisely timing biological processes to occur at the most appropriate times and thereby maximizing resource allocation (12). An important modality through which the circadian clock delivers time-of-day information to output signaling pathways is gating. Circadian gating entails the circadian clock to adjust the sensitivity of output pathways to external and internal stimuli so that the magnitude of the response to a given signal will vary depending on the time of the day. It is assumed that this helps plants filter whether an ambient fluctuation is relevant and ensures that the elicited response is appropriate for the time of the day (12, 13).

In the case of the response to shade, the circadian clock gates it to be maximal at dusk (14) and it was shown that, for plants grown under light/dark cycles, daily shade events were only effective in the promotion of hypocotyl elongation when occurring at the end of the light period (15). Several clock components have been implicated in the regulation of shade signaling (14-18), but only loss of function of the core clock genes *TIMING OF CAB EXPRESSION 1* (*TOC1*) (14) and *CIRCADIAN CLOCK ASSOCIATED 1* (*CCA1*) and *LATE ELONGATED HYPOCOTYL* (*LHY*) (15) was shown to impair temporalization of the response as these mutants reacted equally to shade at both dusk and dawn (14, 15). The underlying molecular mechanisms, however, remain to be elucidated. More recently, it was reported that another clock gene, *EARLY FLOWERING 3* (*ELF3*), represses PIF7 at night mediating the gating of the response at this time (17). However, it is unclear what the physiological relevance of such a finding might be, given that under natural conditions shade poses a stress and is sensed during the light period, not in the middle of the night. We have previously shown that the clock component GIGANTEA (GI) restricts growth at dusk and during the early night by affecting PIF expression and function at multiple levels, including transcriptional and posttranslational mechanisms (19). This function of GI proved to be key to regulate growth rhythms and establish the phase of maximal hypocotyl growth at the end of the night period (19, 20). Here, we show that GI is also required to modulate growth in response to environmental changes fine-tuning the magnitude of the response to shade at dusk. GI achieves this through direct interaction with PIF7 and regulation of the responsiveness of its transcriptional targets to shade at dusk. Furthermore, we pinpoint the epidermis as the tissue where GI function is required in this pathway, reinforcing currents models on circadian clock spatial organization and tissue-level specialization for the regulation of output pathways.

## RESULTS AND DISCUSSION

### *GIGANTEA* mutants display shade avoidance syndrome-related traits and are hypersensitive to shade

GI is a regulator of light signaling and growth and, consequently, *gi* mutants display longer hypocotyls under different light conditions (19, 21, 22). In addition to this increase in hypocotyl length, these mutants also present other traits reminiscent of those that appear in response to shade, such as hyponastic leaves (Figure 1A,B). We therefore examined whether the response to shade is altered in this mutant. We observed that *gi-2* mutants are indeed more responsive to shade and display longer hypocotyls when grown under constant shade conditions (low R:FR ratio < 0.7) (Figure 1C).

**Figure 1.**
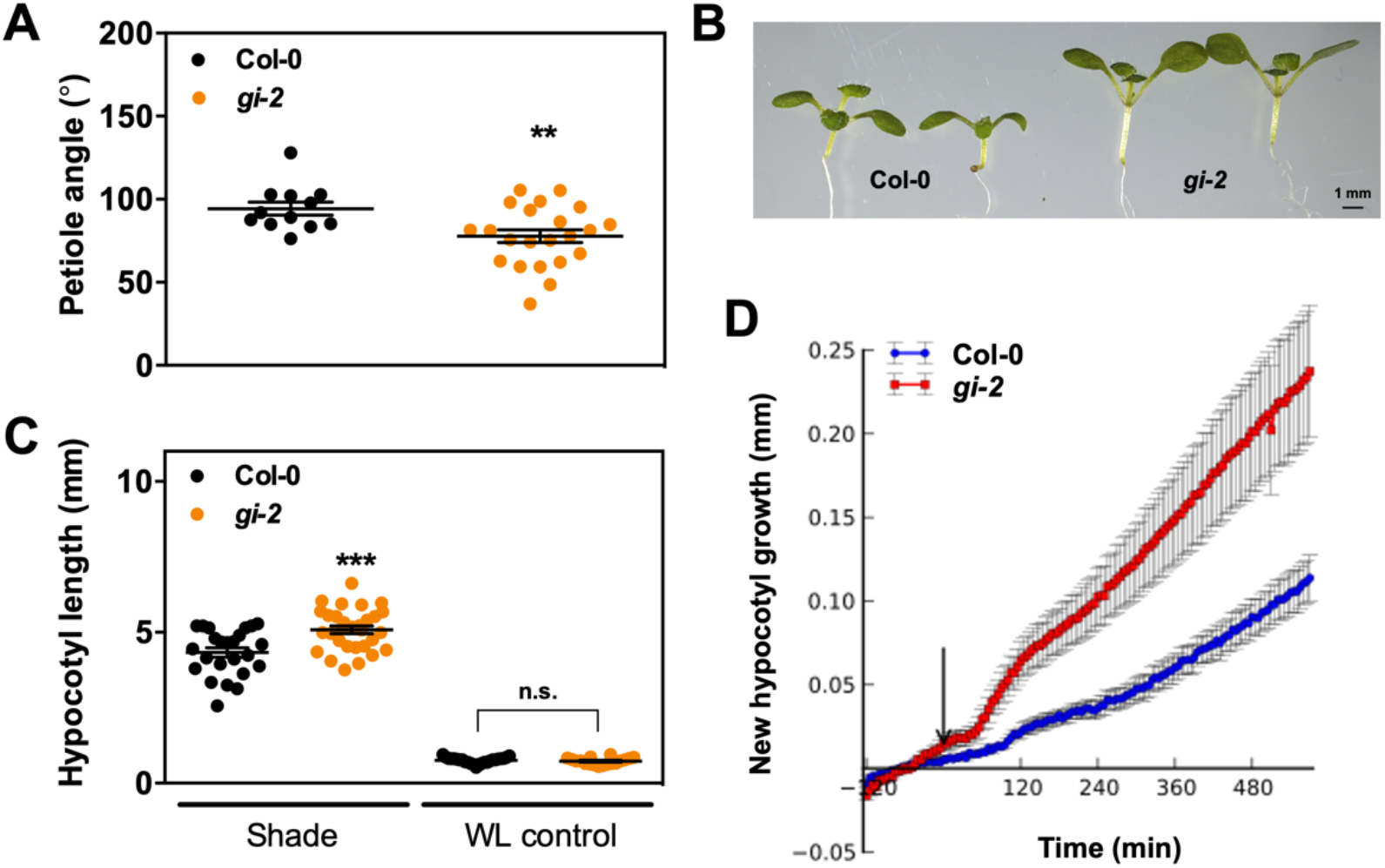
GI is a negative regulator of the response to shade. (*A)* Petiole angle measurements from wildtype (Col-0) and *gi-2* seedlings grown for 10 days under SD conditions. Mean ± SEM, n=12-22; **p<0.01 Student’s *t* test. (*B*) Representative pictures of Col-0 and *gi-2* seedlings grown for 10 days under SDs. (*C*) Hypocotyl length measurements from wildtype (Col-0) and *gi-2* seedlings grown for 4 days under continuous white light and then transferred to constant shade light for 7 days (Shade) or kept in white light (WL control). Mean ± SEM, n=18-30; ***p<0.001, n.s. not significant Tukey’s multiple comparison test. (*D*) New hypocotyl growth observed for Col-0 and *gi-2* seedlings exposed to supplemental FR light to give a R:FR ratio of 0.7. The arrow indicates the start of treatment (t = 0) and hypocotyl growth was monitored for 10h after the treatment. Mean ± SEM, n=12.

In previous studies, it was shown that the fast, initial hypocotyl elongation response to shade is biphasic and occurs after an initial lag phase (23). Examination of this early response to shade revealed that it is also altered in *gi-2* seedlings, which display a shorter lag phase after which elongation occurs in a biphasic mode but at a considerably faster rate compared to wildtype seedlings (Figure 1D). This indicates that *gi-2* mutants not only grow more, but they also respond faster to the change in light quality.

Hormone signaling is central to the regulation of growth-related processes and, consequently, several hormones, including brassinosteriods, gibberellins (GAs) and auxin, play a role in the promotion of hypocotyl elongation in response to shade (24). Auxin, however, seems to be a key player in this pathway (4, 7, 8). With regards to hormone signaling, gene ontology analyses of genes differentially expressed in *gi-2* seedlings have shown that several hormone-related pathways are altered in these mutants, including auxin signaling ((19) and Figure S4A). Additionally, a role for GI in the gating of the response to GAs was recently uncovered and characterized (25). Consequently, we inspected the relevance of both auxin and GAs for the fast response to shade in *gi* mutants (Figure S1). We compared the early response to shade in wildtype (Col-0) and *gi-2* seedlings in the presence or absence of paclobutrazol (PAC, and inhibitor of GA synthesis) and N-1-naphthylphthalamic acid (NPA, an inhibitor of auxin polar transport). Although a significant effect on new hypocotyl growth rate was observed for both treatments, the effect of auxin transport blockage was more drastic, completely nullifying the response (Figure S1). This is consistent with previous observations that auxin plays an essential role in the promotion of hypocotyl elongation in response to shade and indicates that it is required for the function of GI in this pathway.

### GI interacts with PIF7 and functions upstream of it for the regulation of the response to shade

PIF7 plays a prominent role in shade signal transduction as it accumulates in its dephosphorylated (active) form and increases the expression of auxin biosynthetic genes (10, 11). Given that GI interacts with and modulates the activity of several PIFs, we wondered whether interaction with PIF7 could be an underlying mechanism of GI function in the response to shade. We confirmed interaction between GI and PIF7 through several complementary approaches including yeast two-hybrid assays (Figure 2A), *in vitro* pull downs (Figure 2B), and through *in vivo* co-immunoprecipitations (coIPs) in *Arabidopsis thaliana* transgenic lines expressing tagged protein versions of PIF7 and GI (Figure 2C). Genetic analyses also showed that loss of *PIF7* strongly affected hypocotyl elongation specially under shade light and that it completely reduced the long hypocotyl phenotype of *gi-2* in response to shade (Figure 2D). In addition to PIF7, PIF4 and PIF5 have also been implicated in the response to shade (9, 26) and GI is known to modulate the accumulation and activity of these PIFs for the regulation of photoperiodic growth (19). In fact, the rapid response to shade observed in *gi-2* seedlings strongly resembles what occurs under short day (SD) photoperiods, where the expression of growth-promoting genes typically expressed at the end of the night is rapidly induced upon darkness in the absence of GI, resulting in the promotion of hypocotyl elongation at this time (19). We therefore wondered whether GI function in the response to shade occurred mainly through PIF7 or if it included regulation of other PIFs such as PIF4 and PIF5. Hypocotyl length measurements showed that, under SD conditions, it was the loss of *PIF3, PIF5* and both *PIF4* and *PIF5* that had the strongest effect on hypocotyl elongation, strongly reducing the long hypocotyl phenotype of *gi-2* mutants (Figure S1A). Under these conditions, mutations in *PIF7* resulted in a partial suppression of the elongated hypocotyl phenotype of *gi-2* mutants, indicating its partial contribution to this process under this light regime. Under shade light, however, it became evident that *PIF7* played a more prominent role with the *pif7-1* mutation displaying the strongest phenotype and being the one more significantly reducing the hypocotyl elongation phenotype of *gi-2* (Figure S2B). Thus, this genetic interaction supports the hypothesis that *GI* regulates the response to shade upstream of *PIF7*.

**Figure 2.**
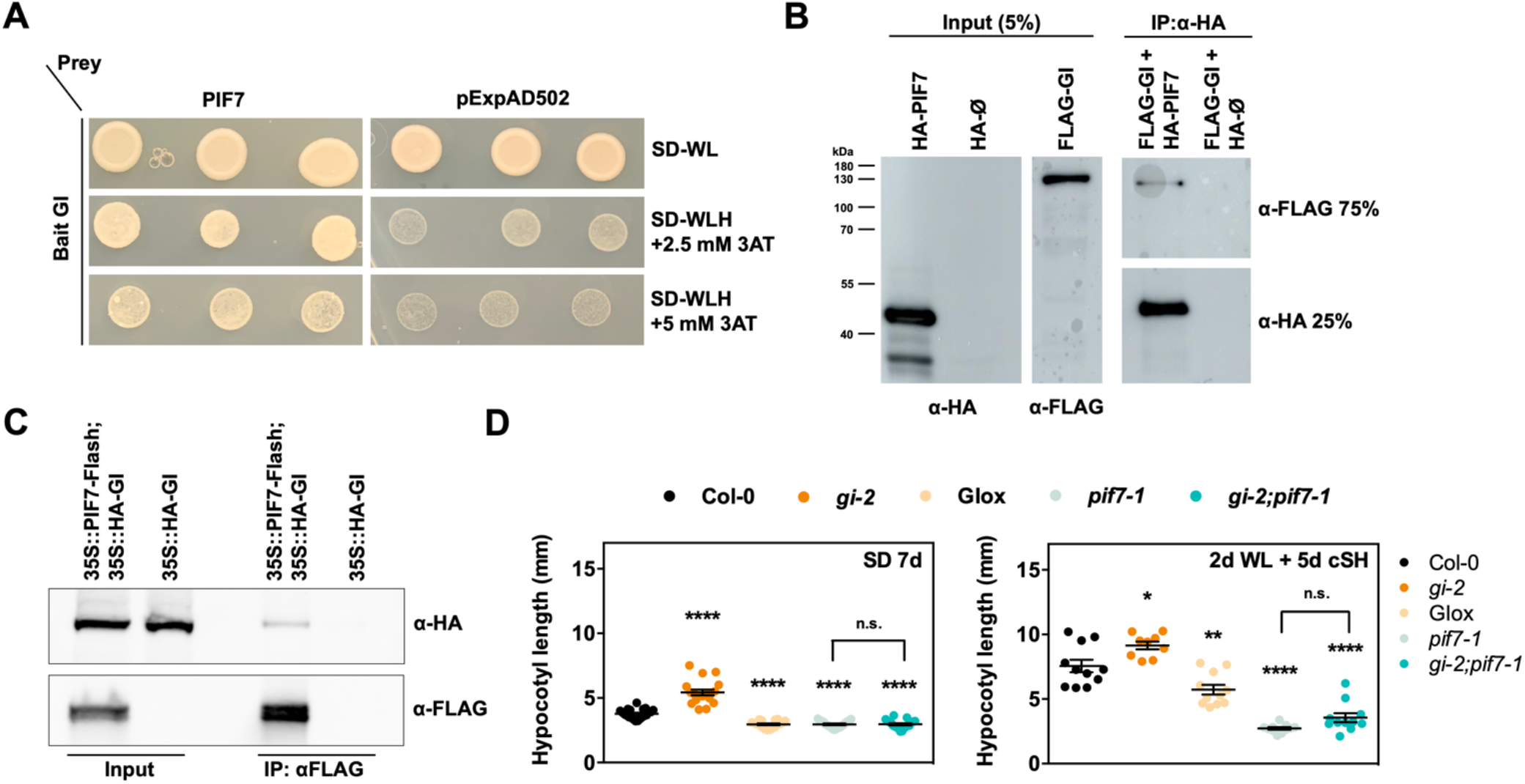
GI interacts with PIF7 and functions upstream of it in the response to shade. (*A)* Yeast two-hybrid (Y2H) assays showing interaction of GI and PIF7 proteins. Bait and prey constructs were co-transformed into yeast cells. SD-WL, minimal medium lacking Trp and Leu; SD-WLH, selective medium lacking Trp, Leu and His, which was supplemented with 2.5 or 5 mM 3AT. (*B*) *In vitro* pull-down assays showing the interaction between GI and PIF7. Proteins were expressed in an *in vitro* transcription and translation system. (*C*) *In vivo* co-immunoprecipitations in *Arabidopsis* transgenic seedlings expressing HA-GI and PIF7-Flash tagged protein versions expressed from the 35S promoter. Seedlings were grown for 7 days under SD conditions and harvested at ZT 8. (*D*) Hypocotyl length measurements from the indicated genotypes grown for 7 days under SD conditions (left panel) or under continuous white light for 2 days and then transferred to constant shade light for 5 days (right panel). Mean ± SEM, n=9-21; ****p<0.0001, ***p<0.001, **p<0.01, *p<0.05, n.s. not significant Tukey’s multiple comparison test.

### GI represses PIF7 activity and affects its binding to target promoter regions in response to shade

GI regulates PIF activity and accumulation through several mechanisms that include transcriptional and post-translational regulation (19, 27). Because *PIF7* expression was observed to be largely unaffected by loss of *GI* function (Figure S3A), we focused on the mechanistic implications of the GI-PIF7 interaction at the protein level. First, we analyzed the effect of GI on PIF7 protein accumulation. We observed that co-infiltration of GI together with PIF7 in transient expression in *Nicotiana benthamiana* leaves did not have any effect on PIF7 accumulation (Figure S3B). A similar trend was observed in Arabidopsis lines expressing a tagged version of PIF7 driven by its own promoter (Figure 3A, S3C,D). These lines behaved like PIF7 overexpression lines (Figure S3D), likely because they were generated in a Col-0 background. Consistent with PIF7 prominent role under shade, we observed that these lines behaved like wildtype under SD but grew more than wildtype seedlings in shade (Figure S3D). Analysis of the accumulation of PIF7 in these lines (in both Col-0 and *gi-2* backgrounds) showed no significant differences in the accumulation of non-phosphorylated (active) PIF7 (under both white light and shade conditions) or in the ratio between its phosphorylated and non-phosphorylated forms (Figure 3A). Hence, GI interaction with PIF7 is unlikely to affect its stability. This finding is not entirely surprising, as previous studies have shown that PIF7 is more stable than other PIFs and, although phosphorylated, it is not rapidly degraded in the light (10, 28). We next performed coIPs in double transgenic lines expressing tagged protein versions of PIF7 and GI driven by endogenous promoter fragments under white light or shade conditions to investigate whether GI preferentially interacts with either the phosphorylated or non-phosphorylated form of PIF7. We found that, although GI is able to interact with both forms of PIF7 under white light, it seems to preferentially interact with the dephosphorylated one in shade (Figure 3B). This could simply be due to increased availability of this form, as it is the one that preferentially accumulates in the nucleus in shade (29). In any case, sequestration of this form, which is the active one, may be a means through which GI interferes with PIF7-mediated expression of shade responsive genes. To further examine the regulation of PIF7 transcriptional activity by GI, we performed transient transcriptional activation assays in *N. benthamiana* leaves using the pPIL1::LUC construct as a reporter of PIF7 transcriptional activity (19, 30). This construct contains the promoter of the well-characterized PIF target gene *PHYTOCHROME INTERACTING FACTOR 3-LIKE 1* (*PIL1*) driving the expression of the firefly luciferase gene (LUC) and it also carries the Renilla luciferase gene (REN) under control of a constitutively expressed promoter as an internal control for normalization. As expected, expression of PIF7 led to an increase in LUC reporter activity (Figure 3C). Co-infiltration of GI together with PIF7, however, led to a significant reduction in pPIL1::LUC activation suggesting that GI interaction with PIF7 is indeed negatively affecting its ability to activate transcription. This observation further supports that GI may negatively regulate the response to low R:FR light by directly interacting with PIF7 and restricting its ability to activate the expression of shade-responsive genes. In fact, expression of one such gene, *YUCCA 8* (*YUC8*), is significantly upregulated in *gi-2* mutants under SD photocycles (Figure S3E). In order to explore the relevance of these findings *in vivo*, we examined the effect of GI on the association of PIF7 to its genomic targets through chromatin immunoprecipitation studies in *A. thaliana* lines. Because PIF7 associates to target sites preferentially under low R:FR light (31), we analyzed the binding of PIF7 to the G-box-containing regions in the promoters of its well-known targets *PIL1* and *YUC8* (10) under these conditions in the presence and absence of GI. For both genes, we observed a significant increase in the enrichment of PIF7 target regions in the immunoprecipitated fractions in the absence of GI (*gi-2* mutant background) (Figure 3D), supporting the notion that GI functions to modulate access of PIF7 to target sites in response to shade. Furthermore, a significant number of genes bound by PIF7 genome-wide (32), including *PIL1* and *YUC8*, are also bound by GI (19) (Figure S3F, hypergeometric test p value < 1.230e-23), and are genes for which the most enriched Gene Ontology (GO) categories are shade avoidance, response to far-red light, and auxin transport (Figure S3G). This suggests that, in addition to sequestration through direct interaction, GI may additionally impede PIF7 binding to target genes similarly to other PIFs (19).

**Figure 3.**
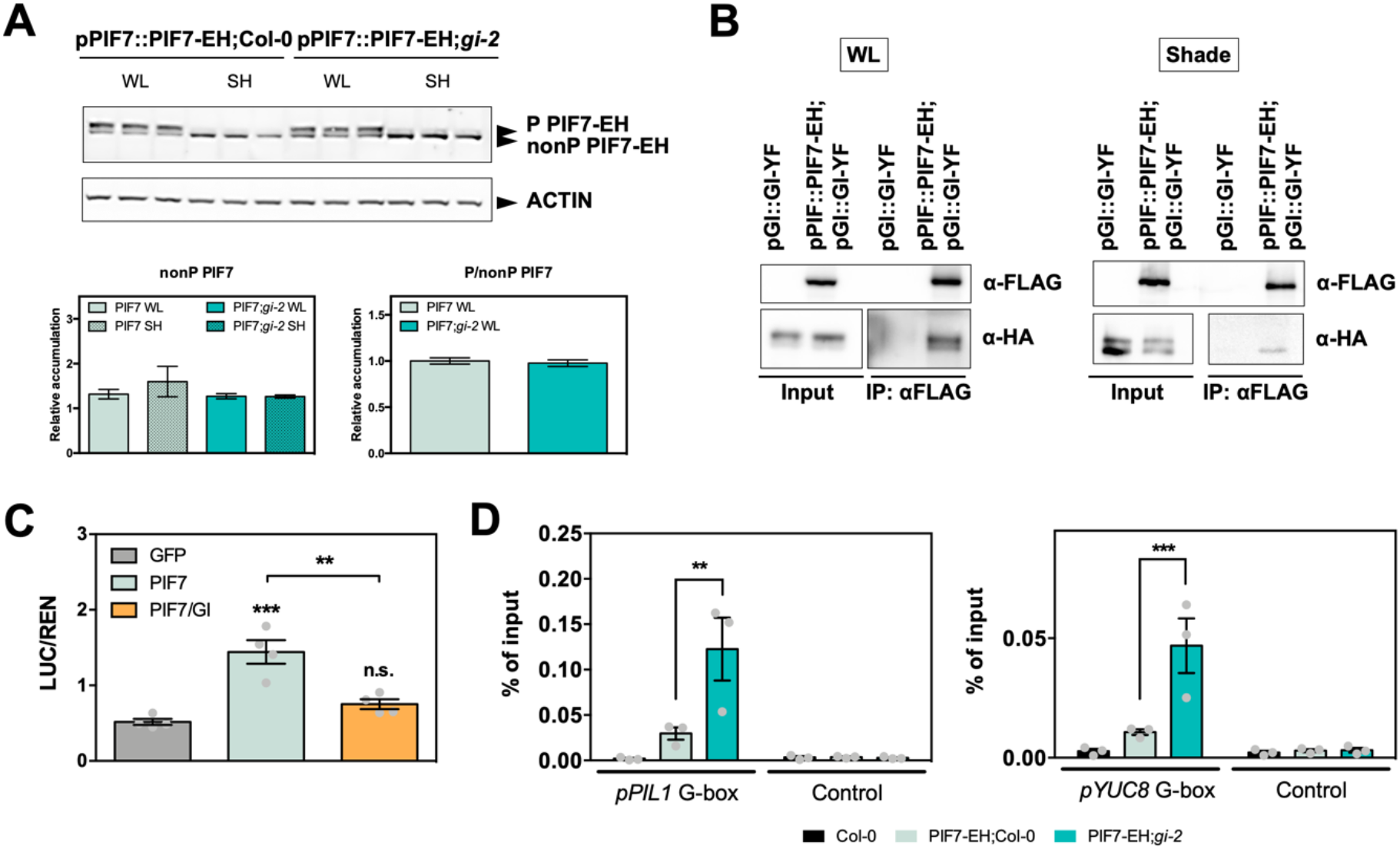
GI interferes with PIF7 transcriptional activity. (*A*) PIF7-ECFP-HA protein accumulation at ZT 9 in the indicated backgrounds grown for 7 d under 10 h light/ 14 h dark conditions. Seedlings were either treated with shade for 1 h prior to harvesting (ZT 8 to 9) or kept in the light. The lower panels show the quantitation of the non-phosphorylated (left) and the ratio between phosphorylated and non-phosphorylated forms (mean ± SEM of 3 biological replicates). Protein levels were normalized against ACTIN levels. (*B*) Co-immunoprecipitation assays in *Arabidopsis* transgenic seedlings expressing PIF7-ECFP-HA and GI-YPET-FLAG from their respective endogenous promoters. Seedlings were grown for 7 d under 10 h light/ 14 h dark conditions and then either treated with shade for 1 h prior to harvesting (ZT 8 to 9) or kept in the light. (*C*) Transactivation assays in *N. benthamiana* leaves. Different effectors were co-expressed with the *pPIL1*::LUC reporter construct. Luminescence was measured 3 days post-infiltration and the ratio LUC/REN was calculated. Results show mean ± SEM (n=4). ***p<0.001, **p<0.01, n.s. not significant Tukey’s multiple comparison test. (*D*) ChIP assays of 7-day-old seedlings grown under 10 h light/ 14 h dark conditions and then treated with shade for 1 h prior to harvesting (ZT 8 to 9). The enrichment of the specified regions in the immunoprecipitated samples was quantified by qPCR. Values represent mean ± SEM of 3 biological replicates. ***p<0.001, **p<0.01 Tukey’s multiple comparison test.

### GI restricts the shade-responsive expression of PIF7 target genes

Considering our findings on the modulation of PIF7 activity by GI, we next investigated the relevance of the interaction between GI and PIF7 for the expression of shade-responsive genes genome-wide. To this end, we performed an RNA sequencing (RNAseq) experiment and identified differentially expressed genes (DEGs) in *gi-2* compared to wildtype (Col-0) plants. Comparison of this set of genes with a set of PIF7-dependent genes (DEGs in *pif7-1*, as defined by Chung and coworkers (32)) evidenced a significant overlap (hypergeometric test p value < 1.611e-13) where the expression of a third of PIF7-dependent genes is mis-regulated as a consequence of *GI* loss of function (Figure 4A). Importantly, half of these shared target genes are bound by GI at their promoter regions. In terms of GO enrichment, shared target genes were again found to be mainly involved in growth-related processes, auxin signaling, and the response to far-red light (Figure S4A) and include well-known PIF7 target genes such as *PIL1, YUC8, ARABIDOPSIS THALIANA HOMEOBOX PROTEIN 2* (*ATHB2*), and *PHYTOCHROME RAPIDLY REGULATED 1* (*PAR1*) (10, 33). These are genes typically induced upon shade and, consistent with the repressive effect of GI on PIF7 transcriptional activity, they appeared to be significantly upregulated in *gi-2* (Figure 4B). Given the effect of GI on the hypocotyl elongation response to shade, we next investigated whether the shade-promoted induction of their expression was also affected in *gi-2*. To this end, we grew Arabidopsis seedlings under light/dark photocycles (10 h light/ 14 h darkness) and performed a 1h long shade treatment around dusk (Zeitgeber Time (ZT) 8). This is the time when Arabidopsis is more responsive to shade (15) and GI is more highly expressed (19, 34, 35). We observed that, indeed, these genes are more strongly induced by shade in *gi-2* and that this induction is dependent on PIF7 function (Figure 4C, S4B). Interestingly, these genes were still slightly induced in *gi-2;pif7-1* mutants, most likely due to the participation of other factors controlled by GI in their regulation upon shade, such as PIF4 and 5 (18, 19). Noteworthy, the expression of neither *PIF7* nor *GI* was induced, further supporting the notion that the induction of PIF7 target genes is the consequence of a mechanism operating post-transcriptionally (Figure 4C, S4B). Altogether, our findings point to a function of GI in modulating the magnitude of the response to shade by regulating PIF7-mediated transcriptional activation.

**Figure 4.**
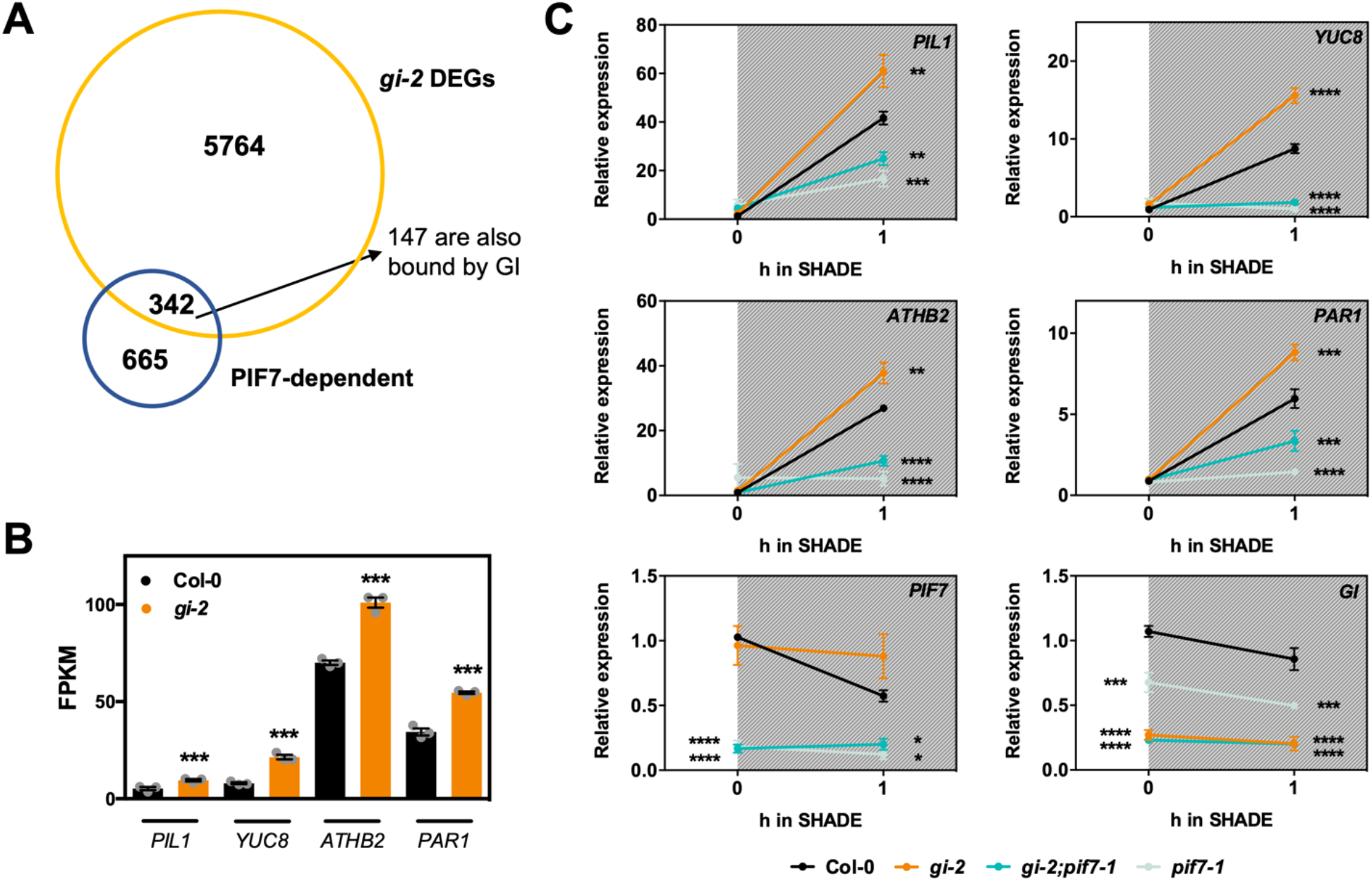
GI restricts the shade induced expression of PIF7 target genes. Overlap between DEGs in *gi-2* and a set of PIF7-dependent genes (32) (intersection p value < 1.611e-13). Expression levels in FPKM of several PIF7 targets genes as identified by RNAseq (***p<0.001). (*C*) Relative expression of *PIL1, YUC8, ATHB2, PAR1, PIF7* and *GI* in the indicated backgrounds in response to a 1 h long treatment with shade at dusk (ZT 8 to 9). Seedlings were grown for 7 days under 10 h light/ 14 h dark photocycles. Mean ± SEM of 3 biological replicates. White and gray shadings represent light and shade periods, respectively. ****p<0.0001, ***p<0.001, **p<0.01, *p<0.05 Tukey’s multiple comparison test.

### GI gates the sensitivity to shade at dusk and functions in the epidermis to regulate shade signaling

It has been previously shown that the response to shade is maximal around dusk and a role for light and circadian signaling components was proposed (14, 15, 17). Hence, we next examined the physiological relevance of GI function in shade signaling and evaluated its role in the gating of the response to shade. Consistent with previous observations, shade events occurring in the morning were ineffective, while shade perception in the afternoon had a significant effect on hypocotyl elongation (Figure S5A). In this context, we observed that *gi-2* mutants were hypersensitive to shade at dusk and grew significantly more than Col-0 seedlings. Moreover, this phenotype was observed to be dependent on PIF7 function (Figure S5A). In order to take a closer look at the response at dusk, we performed 2 h shade treatments at different ZTs around dusk time (ZT 6, 7, 8, 9, and 10; seedlings grown under 10 h light/ 14 h darkness photocycles) and quantified the effect of the treatment on hypocotyl elongation in *gi-2* compared to Col-0 seedlings (Figure 5A). These experiments allowed us to confirm that GI is required to modulate the magnitude of the response to shade and that its function is more relevant at dusk, at ZT 8 to 10 (Figure 5A).

**Figure 5.**
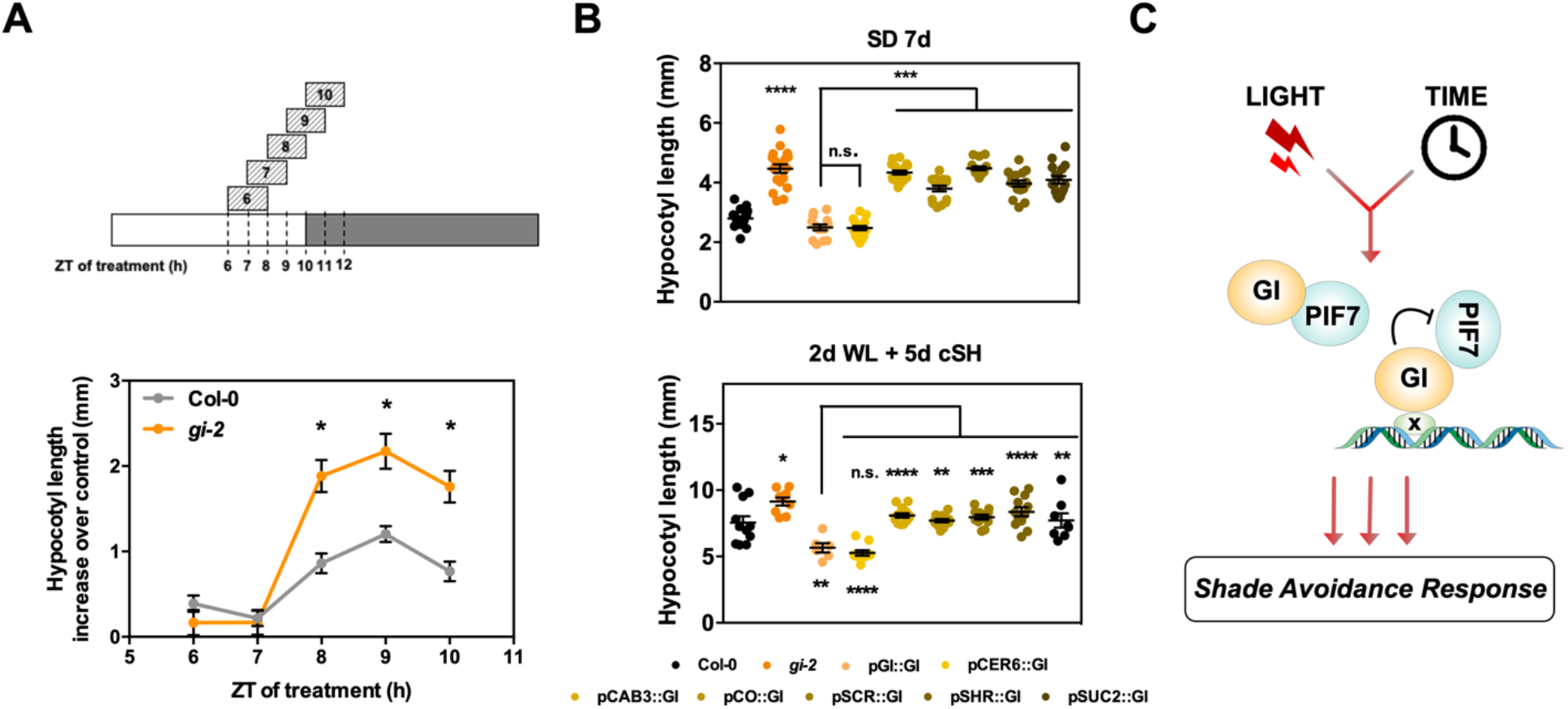
GI functions in the epidermis to modulate the response to shade at dusk. (*A*) Hypocotyl length increase (measured as the difference between shade treated and white light kept seedlings) of seedlings grown for 3 days under 10 h light/ 14 h dark conditions and either exposed every day to shade for 2 h at the times indicated or kept in the light (mean ± SEM, n=15-23) (*p<0.05 Tukey’s multiple comparison test). A scheme of the experimental set up is shown in the upper panel. (*B*) Hypocotyl length measurements from the indicated genotypes grown for 7 days under SD conditions (upper panel) or under continuous white light for 2 days and then transferred to constant shade light for 5 days (lower panel). Mean ± SEM, n=14-21; ****p<0.0001, ***p<0.001, **p<0.01, *p<0.05, n.s. not significant Tukey’s multiple comparison test. (*C*) Model depicting GI function in the response to shade at dusk. At this time, GI-mediated repression of PIF7 (through direct interaction and likely also through occupation of PIF7 genomic targets) provides a means through which the clock controls the magnitude of the response to the changes in light quality.

In Arabidopsis seedlings, shade sensed in the cotyledons triggers a transcriptional response to promote auxin biosynthesis, which then travels to the hypocotyl to induce cell elongation in response to the perceived changes in light quality (36, 37). For this, the epidermis was shown to play an important role, as in this tissue auxin functions, at least partially, to induce brassinosteroid-mediated cell elongation (38). In recent years, it is becoming evident that the circadian system in Arabidopsis is spatially organized, with circadian clocks differentially processing specific environmental signals in different tissues to coordinate individual physiological responses (39, 40). To gain insights into the spatial characteristics of GI-mediated modulation of the response to shade, we investigated the effect of tissue-specific (TS) expression of GI on hypocotyl elongation under both SD photoperiods and in response to low R:FR ratio. To this end, we transformed *gi-2* null mutants with a suite of constructs in which the coding sequence of *GI* is expressed from an endogenous promoter fragment (pGI) or from different TS promoters (38, 41) and analyzed their ability to complement the mutant phenotype in terms of hypocotyl elongation. The TS promoters used comprised pCAB3 (mesophyll), pCER6 (epidermis), pCO2 (cortex), pSCR (endodermis), pSHR (stele) and pSUC2 (phloem companion cells) (38, 41). As expected, we observed that *GI* expressed from its endogenous promoter rescued the long hypocotyl phenotype of *gi-2* in SD photoperiods and in response to shade (Figure 5B). Noteworthy, from all TS promoters tested, only *GI* expressed in the epidermis rescued the long hypocotyl phenotype of *gi-2* similarly to pGI (Figure 5B), even though *GI* was expressed at considerable levels in all lines (Figure S5B). Hence, GI function is required in the epidermis to modulate hypocotyl elongation under varying light conditions, which is in line with previous observations about TS functions of the clock which showed that the clock in the epidermis is essential for cell elongation in response to temperature (40).

Altogether, our findings support a model (Figure 5C) where the circadian clock, through the function of GI, fine-tunes the magnitude of the response to changes in light quality at dusk by directly impinging on key transcriptional regulators of the shade signaling pathway and highlights the relevance of TS circadian function for the regulation of specific output processes.

## CONCLUDING REMARKS

The circadian clock is a complex molecular network that confers plants (and other organisms) the ability to phase biological processes to the most appropriate time of the day and year in resonance with the environment. Importantly, it also modulates the sensitivity of specific signaling pathways to internal and external cues at particular phases. The ability of the circadian network to integrate multiple signals together with its robust rhythmicity is thought to help plants discern noise from the key environmental signals, thereby reducing the effects of non-informative environmental variability, such as stochastic variation in the daily light intensity (42).

Like many other physiological processes, the response to shade is under circadian control and, in terms of phasing, the clock seems to function to temporalize the sensitivity to shade signals gating it towards dusk (14, 15). Although several clock components have been shown to affect the SAR (16-18), how gating of the response is achieved at dusk time was poorly understood at the molecular level. Here, we provide evidence on a mechanistic link between the central oscillator and the response to shade and show how it functions specifically in the epidermis to modulate the response to shade at dusk. Hence, our work uncovers an important mechanism by which a tissue-specific circadian oscillator modulates plastic growth in dynamic environments.

## MATERIALS AND METHODS

### Plant material and growth conditions

Wildtype, mutant, and transgenic lines used in this study were *Arabidopsis thaliana* ecotype Columbia 0 (Col-0). *gi-2* (34), 35S::HA-GI;*gi-2* (35), GIox (19), pGI::GI-YPET-FLAG;*gi-2* (19), 35S::PIF7-Flash;*pif7-2* (10), *pif7-1* (SALK_062756/ SALK_037763) (28), *pif7-2* (Syngenta collection of sequenced T-DNA insertional mutants, line 622) (28), *pif3-1* (SALK_030753) (43), *pif4-101* (Garlic_114_G06) (9), *pif5-1* (SALK_087012) (44), *gi-2;pif3-1* (19), *gi-2;pif5-1* (19), and *gi-2; pif4-101;pif5-1* (19) have been previously described.

Seeds were chlorine gas sterilized and plated on 0.5x Murashige and Skoog medium (MS, Caisson Laboratories) with 0.8% agar (Sigma). After stratification in the dark at 4 °C for 3 days, plates were transferred to a Percival incubator (Percival-scientific.com) set to the indicated light conditions with light supplied at 80 μmol m^-2^ s^-1^ by cool-white fluorescent bulbs and a constant temperature of 22 °C.

For shade light treatments of seedlings grown on plates, they were transferred to a Percival incubator provided with LEDs set to supply shade-simulating light (red light 13 µmol m^2^s^-1^; far-red light 20.2 µmol m^2^s^-1^; blue light 1.23 µmol m^2^s^-1^; R:FR 0.7).

### Generation of transgenic lines

Transgenic pPIF7::PIF7-ECFP-HA and tissue-specific GI lines were generated by Agrobacterium-mediated floral dip transformation of Col-0 and *gi-2* plants, respectively. To this purpose, *Agrobacterium tumefaciens* strain GV3101 was transformed with the binary vectors pH-pPIF7::PIF7-ECFP-HA, pB-pCAB3::GI-YPET-Flag, pB-pCER6::GI-YPET-Flag, pB-pCO2::GI-YPET-Flag, pB-pSCR::GI-YPET-Flag, pB-pSHR::GI-YPET-Flag, and pB-pSUC2::GI-YPET-Flag (described below), respectively. Lines with single insertions were selected scoring for a 3:1 survival ratio in growth medium supplied with the appropriate antibiotic in the F2 progeny of individual F1 plants and were brought to homozygosity in the F3 generation.

### Construction of binary vectors

The full-length cDNA encoding PIF7 (without the stop codon) was amplified by PCR and cloned into the pDONR207 vector (Invitrogen) (primers used are listed in Supplementary Table S1). pENTR-GI-stop has been described earlier (45). In the case of the endogenous *PIF7* promoter sequence, 2000 bp upstream of the *PIF7* start codon were amplified by PCR using the primers listed in Supplementary Table S1 and cloned into the pDONR P4-P1R vector (Invitrogen) by Gateway BP recombination reaction (Invitrogen). The pDONR P4-P1R vectors containing the different tissue-specific promoters have already been described (38, 41), as have the pDONR P2R-P3 vectors containing the ECFP-HA and YPET-FLAG tags (19). Selected promoter, gene, and fluorescent tag combinations were finally cloned by MultiSite Gateway reaction (Invitrogen) into either pH7m34GW (pPIF7::PIF7-ECFP-HA) or pB7m34GW (pB-pCAB3::GI-YPET-Flag, pB-pCER6::GI-YPET-Flag, pB-pCO2::GI-YPET-Flag, pB-pSCR::GI-YPET-Flag, pB-pSHR::GI-YPET-Flag, and pB-pSUC2::GI-YPET-Flag) binary destination vectors from the University of Ghent collection (https://gateway.psb.ugent.be).

To perform protein stability and transactivation assays in transient expression in *N. benthamiana*, the PIF7 coding sequence was subsequently transferred from the pDONR207 to the pEarleyGate201 (46) binary destination vector by Gateway LR recombination reaction (Invitrogen). pEarleyGate201-GFP and pEarleyGate202-GI vectors have already been described (19), as has the pPIL1::LUC construct (19).

### Yeast two-hybrid analyses

For yeast two-hybrid analyses, the Clontech matchmaker GAL4 System was used according to the manufacturer’s instructions. The CDSs of GI and PIF7 were cloned into Gateway compatible vectors, obtained by respective modification of the pGBKT7 AND pGADT7 vectors (Clontech). Specifically, pDONR207-PIF7 was recombined with pGADT7-GW using the Gateway LR recombination reaction (Invitrogen) to produce PIF7 fused to the Gal4-activation domain (AD) and GI was introduced from the pENTR-GI vector described above into pGBKT7-GW to fuse with the GAL4 DNA-binding domain. Constructs were co-transformed into the yeast strain AH109 and grown at 28ºC on SD (selective drop out) agar medium lacking tryptophan (Trp) and leucine (Leu) (SD-WL). Protein interactions were assayed by the nutritional requirement on histidine (His) on plates lacking Trp, Leu and His (SD-WLH). SD-WLH plates were supplemented with either 2.5 or 5 mM 3-amino-1,2,4-triazole (3-AT, Sigma-Aldrich).

### *In vitro* pull-downs

To express proteins in the cell-free system, all inserts were transferred by Gateway LR recombination reaction (Invitrogen) into Gateway compatible modified pTnT vectors (Promega) containing an N-terminal HA or FLAG tag as specified in each case. pDONR207-PIF7 was recombined with the pTnT-HA vector to produce PIF7 fused to HA. The construct containing GI in the pTnT-FLAG vector has already been described (19). For the *in vitro* pull-down assays, proteins were co-expressed using TnT® SP6 High-Yield Wheat Germ Protein Expression System (Promega) as per manufacturer’s instructions. Five percent of the reactions (2.5 µl) were used to verify expression of the proteins (input) and the remaining extract was immunoprecipitated as earlier described (47) using a modified IP buffer (50 mM Tris-HCl pH 7.6, 150 mM NaCl, 0.5% Triton X-100, 1x protease inhibitor cocktail (Roche), 1x Phosphatase Inhibitors I&II (Sigma) and 25 μM MG-132) and the anti-HA 3F10 antibody (Roche).

### Transient expression in *Nicotiana benthamiana*

*Agrobacterium tumefaciens* strain GV3101 was used in all instances. In the case of the pPIL1::LUC construct, it additionally contained the pSoup helper plasmid. *A. tumefaciens* cells containing the respective constructs and the p19 silencing suppressor were grown overnight at 28 °C in liquid LB medium supplemented with the appropriate antibiotics. Cultures were pelleted, resuspended in 10 mM MES-KOH pH 5.6, 10 mM MgCl_2,_ 150 µM acetosyringone to a final OD_600_ of 0.5, and incubated for 2h at room temperature. The suspensions were then mixed and infiltrated in *N. benthamiana* leaves at a final OD_600_=0.1 each, except for p19 which was infiltrated at a final OD_600_=0.05. Samples were harvested 3 days post-inoculation.

### Protein immunoprecipitation

Approximately 1 g of 10-day-old *Arabidopsis* seedlings grown in SDs were harvested at the times indicated and frozen in liquid nitrogen. Immunoprecipitations were performed as earlier described (19) with the following modifications. Samples were ground with mortar and pestle in liquid nitrogen and resuspended in 2 ml of IP buffer (50 mM Tris-HCl pH 7.6, 150 mM NaCl, 5 mM MgCl_2_, 0.1% NP-40, 10% glycerol, 2 mM PMSF, 1x protease inhibitor cocktail (Roche), 1x Phosphatase Inhibitors I&II (Sigma) and 50 μM MG-132). Extracts were transferred to a dounce tissue grinder and homogenized before being clarified twice by centrifugation at 4 °C. Total protein concentration was quantified by DC Protein Assay (Bio-Rad) and normalized to 1.875 mg/ml. Three percent of the extracts was used to verify proteins levels (input). For immunoprecipitations, extracts were incubated with anti-Flag M2 antibody (Sigma) antibody for 2 h with gentle rotation at 4 °C. Subsequently, 25 µl of magnetic protein G Dynabeads (Invitrogen) pre-washed with IP buffer were added to the samples and incubated for 2 h with gentle rotation at 4 °C. The samples were finally washed 3x with IP buffer and the precipitated protein was eluted by heating beads at 95 °C for 5 min in 40 μl of 2x SDS-PAGE loading buffer. 10 and 30 µl of the eluate were separately analyzed by Western blot to detect the immunoprecipitated and co-immunoprecipitated proteins, respectively.

### Protein accumulation analyses

To determine protein levels in transient experiments in *N. benthamiana*, samples were homogenized with 3 volumes of 2x SDS-PAGE loading buffer and boiled at 95 °C for 5 min. Samples were then clarified by centrifugation at room temperature and analyzed by Western blot. For normalization, GFP-HA was used as internal loading control.

For the analysis of PIF7-ECFP-HA protein levels in *Arabidopsis* seedlings, samples were homogenized with 100 µl of extraction buffer (50 mM Tris-HCl pH 7.6, 150 mM NaCl, 5 mM MgCl_2_, 0.1% NP-40, 10% glycerol, 2 mM PMSF, 1x protease inhibitor cocktail (Roche), 1x Phosphatase Inhibitors I&II (Sigma) and 50 μM MG-132) and clarified twice by centrifugation at 4 °C. Total protein concentration was quantified by DC Protein Assay (Bio-Rad) and 40 ug of each sample was subsequently analyzed by Western blot. ACTIN levels in the samples were used for normalization.

### Western blot detection and quantitation

Protein extracts in SDS-PAGE loading buffer were boiled at 95 °C for 5 min and separated in 4-15 % SDS-PAGE gels. Proteins were then transferred to PVDF membranes (Amersham), which were then stained with Ponceau S to assess transfer and loading. Finally, membranes were cut and immunodetected with either Horse Radish Peroxidase (HRP)-conjugated 3F10 anti-HA (1:2000, Roche) or HRP-conjugated Flag M2 (1:2000, Sigma) antibodies (1:2000, Roche). The ACTIN loading control was detected using anti-ACTIN C4 mouse antibody (1:500, Millipore) followed by incubation with NA931V HRP-conjugated anti-mouse secondary antibody (1:10000, Sigma). Chemiluminescence was detected with the Supersignal West Pico, Dura, and Atto substrates (Thermo Scientific) and imaged with an ImageQuant 800 system (Amersham). Protein levels were quantified using NIH ImageJ software (https://imagej.nih.gov/ij/).

### Transactivation assays

Reporter and effector constructs were co-infiltrated in *N. benthamiana* leaves and 3 days post infiltration (dpi) trancriptional activation assays were performed. To this end, firefly (LUC) and the control *Renilla* (REN) luciferase activities were assayed from leaf extracts with the Dual-Glo Luciferase Assay System (Promega) and quantified with a GloMax-Multi Detection System (Promega).

### RNA extraction and qRT-PCR

Total RNA was isolated with the GeneJET Plant RNA Purification Kit (Thermo Scientific). For cDNA synthesis, 1 μg of total RNA was digested with DNase I (Roche) and reverse-transcribed using the NZY First-Strand cDNA Synthesis Kit (nzytech). Synthesized cDNA was amplified by real-time quantitative PCR (qPCR) with TB Green Premix Ex Taq (Tli RNaseH Plus) (Takara) using the QuantStudio 3 system (Applied Biosystems). *PROTEIN PHOSPHATASE 2A* (*PP2A*) (AT1G13320) was used as the normalization control in Figure S3A and E and *ISOPENTENYL PYROPHOSPHATE:DIMETHYLALLYL PYROPHOSPHATE ISOMERASE 2* (*IPP2*) (AT3G02780) was used in Figure 4C and S4B and 5B. Primer sequences are listed in Supplementary Table S1.

### Chromatin immunoprecipitation (ChIP)

ChIP assays were performed as earlier described (19) with no modifications, except that 2 g of fresh weight whole seedlings were harvested for each genotype (3 biological replicates each) and 1.5 ug of anti-HA 3F10 (Roche) antibody were used for immunoprecipitation. Immunoprecipitad DNA was analyzed by qPCR and reactions were performed in triplicate with TB Green Premix Ex Taq (Tli RNaseH Plus) (Takara) using the QuantStudio 3 system (Applied Biosystems). Primers used were as in previous studies (10) and are listed in Supplementary Table S1. Enrichment in each sample was calculated relative to the input and expressed in percentage.

### RNA sequencing

Three biological replicates of each wildtype (Col-0) and *gi-2* seedlings grown under SD conditions for 7 days. were collected at ZT 0 (whole seedlings) and snap-frozen in liquid nitrogen. Total RNA was isolated using the GeneJET Plant RNA Purification Kit (Thermo Scientific) and digested with DNase Max (Qiagen). Preparation of RNA library (non-directional 250∼300 bp insert cDNA library (NEB)) and transcriptome sequencing (Illumina Novaseq 6000 paired-end 150) was conducted by Novogene Corporation Inc. (Davis Research Lab, USA). Raw reads were aligned to the TAIR10 genome using STAR (v2.5.5) (48) with the method of Maximal Mappable Prefix (MMP). Mapped reads were counted by HTSeq v0.6.1 and then FPKM (Reads Per Kilobase of exon model per Million mapped reads) of each gene was calculated based on the length of the gene and reads count mapped to this gene. Differential expression was performed using the DESeq2 R package (2_1.6.3) (49). Genes with adjusted p-value < 0.05 and |log2(FoldChange)| > 0 were considered as differentially expressed.

### Genome-wide data set analyses

All datasets used for genome-wide gene expression analysis are compiled in the datasets Supplementary File. The statistical significance of the overlap between gene lists was calculated using the hypergeometric test.

For the identification of over-represented Gene Ontology (GO) categories in gene lists, the web-based tool PANTHER (http://www.pantherdb.org/) (50) was used. For the significantly enriched categories (p-value<0.05) in each subset, enrichment scores were calculated as -log_10_ (p), where p is the p-value from the enrichment analysis.

### Physiological measurements

To analyze hypocotyl length, evenly spaced seedlings were grown on plates under the indicated light conditions and photoperiod. At the specified time, seedlings were scanned and images were analyzed using NIH ImageJ software (https://imagej.nih.gov/ij/).

Dynamic hypocotyl elongation was quantified using time-lapse imaging as previously described (23) with some modifications. We adapted an automated rotation stage (Griffin Motion, Holly Springs, NC) to rotate seedlings grown on folded pieces of whatman filter paper soaked with 0.5x Murashige and Skoog medium (with 0.8% Agar), in 1.1 mL cylindrical plastic tubes (Thermo Scientific). Seedlings were grown in a fluorescent white light growth chamber (Percival Scientific) under long day (16 hr light/8 hr dark) at 22°C for 5 days. 4 hours prior to dusk on the 5^th^ day, seedlings were transferred to an LED chamber (Percival Scientific), and imaged for 12 hours. LED-simulated white light and LED-shade (low R/FR) conditions were as described previously (23). For drug treatments, seedlings were immersed in 500 μL of the indicated concentration of NPA or PAC 12 hours prior to imaging. The liquid was allowed to soak the seedlings for 2 hours, then aspirated. Seedlings were then air-dried (while still on wet MS-soaked whatman paper) for 10 hours prior to imaging. Shade treatment was applied as indicated. Hypocotyl length was quantified using the HyDE algorithm, as previously described (23).

The analyses on the gating of the response to shade were performed as previously described (15). Briefly, evenly spaced seedlings were grown on plates under 10 h light/ 14 h dark photoperiod. Each day at the specified times (for three consecutive days), seedlings were transferred to a Percival incubator provided with LEDs set to supply shade-simulating light (as specified above) or kept in the control conditions (no shade treatment). After the third day, seedlings were scanned and images were analyzed using NIH ImageJ software (https://imagej.nih.gov/ij/). Hypocotyl length increase upon shade treatment at the different ZTs was calculated for shade-treated plants relative to non-treated plants.

## ACKNOWLEDGEMENTS

We thank M. Blázquez for critical reading of the manuscript. Work in the authors’ laboratories has been funded by the Spanish Agencia Estatal de Investigación and Generalitat Valenciana (grants PID2020-119491GA-I00 and CIDEGENT/2020/037 to M.A.N.) and the National Institute of General Medical Sciences of the National Institutes of Health (under award numbers R37 GM067837 to S.A.K.). J.C. is an investigator from the Howard Hughes Medical Institute.

