## Supplemental Figures for "Epidermal GIGANTEA adjusts the response to shade at dusk by directly impinging on PHYTOCHROME INTERACTING FACTOR 7 function"

SUPPEMENTARY FIGURES

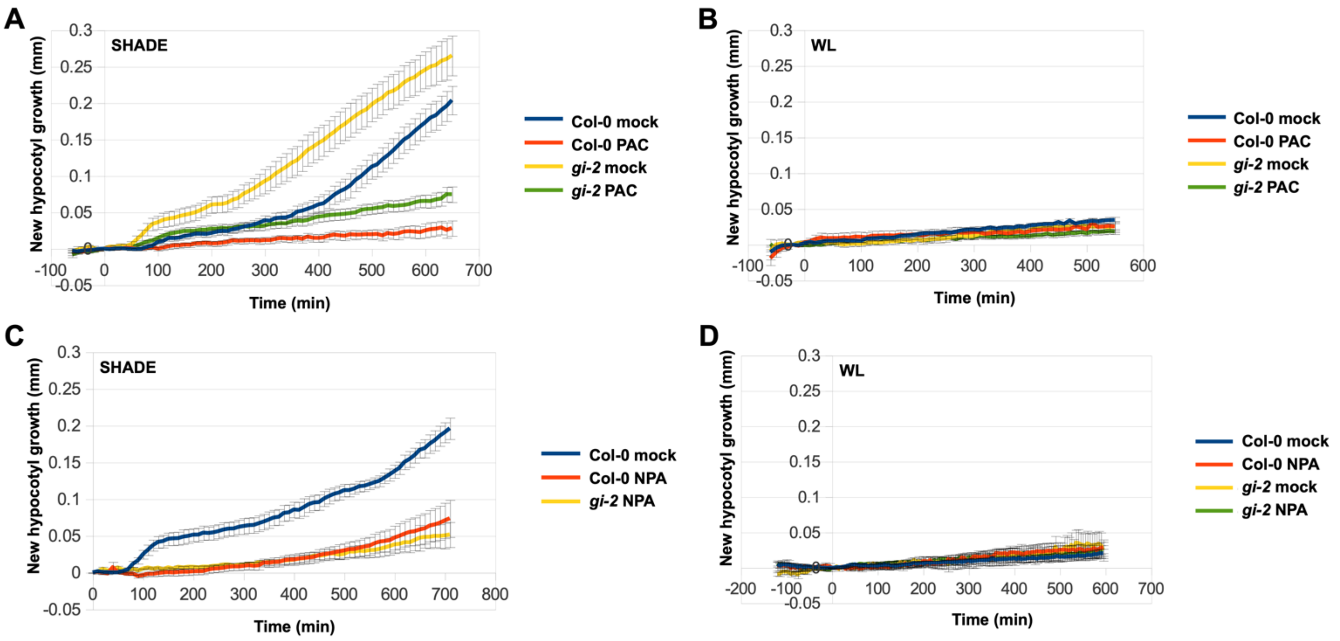


**Figure S1.**

(*A*, *B*) New hypocotyl growth measured for Col-0 and *gi-2* seedlings grown in the presence (PAC) or absence (mock) of paclobutrazol exposed to supplemental FR light to give a R:FR ratio of 0.7 (*A*) or kept in white light (*B*). The start of treatment is t = 0. Mean ± SEM, n=12. (*C*, *D*) New hypocotyl growth measured for Col-0 and *gi-2* seedlings grown in the presence (NPA) or absence (mock) of N-1-naphthylphthalamic acid exposed to supplemental FR light to give a R:FR ratio of 0.7 (*C*) or kept in white light (*D*). The start of treatment is t = 0. Mean ± SEM, n=12.

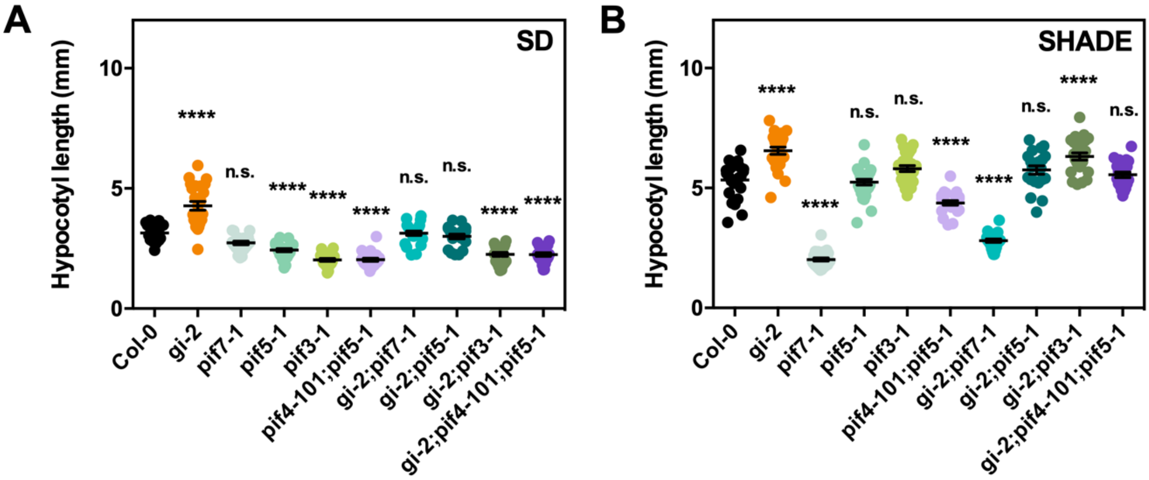


**Figure S2.**

(*A*, *B*) Hypocotyl length measurements from the indicated genotypes grown for 7 days under SD conditions (*A*) or under continuous white light for 2 days and then transferred to constant shade light for 5 days (*B*). Mean ± SEM, n=19-28; ****p<0.0001, ***p<0.001, **p<0.01, *p<0.05, n.s. not significant Tukey's multiple comparison test.

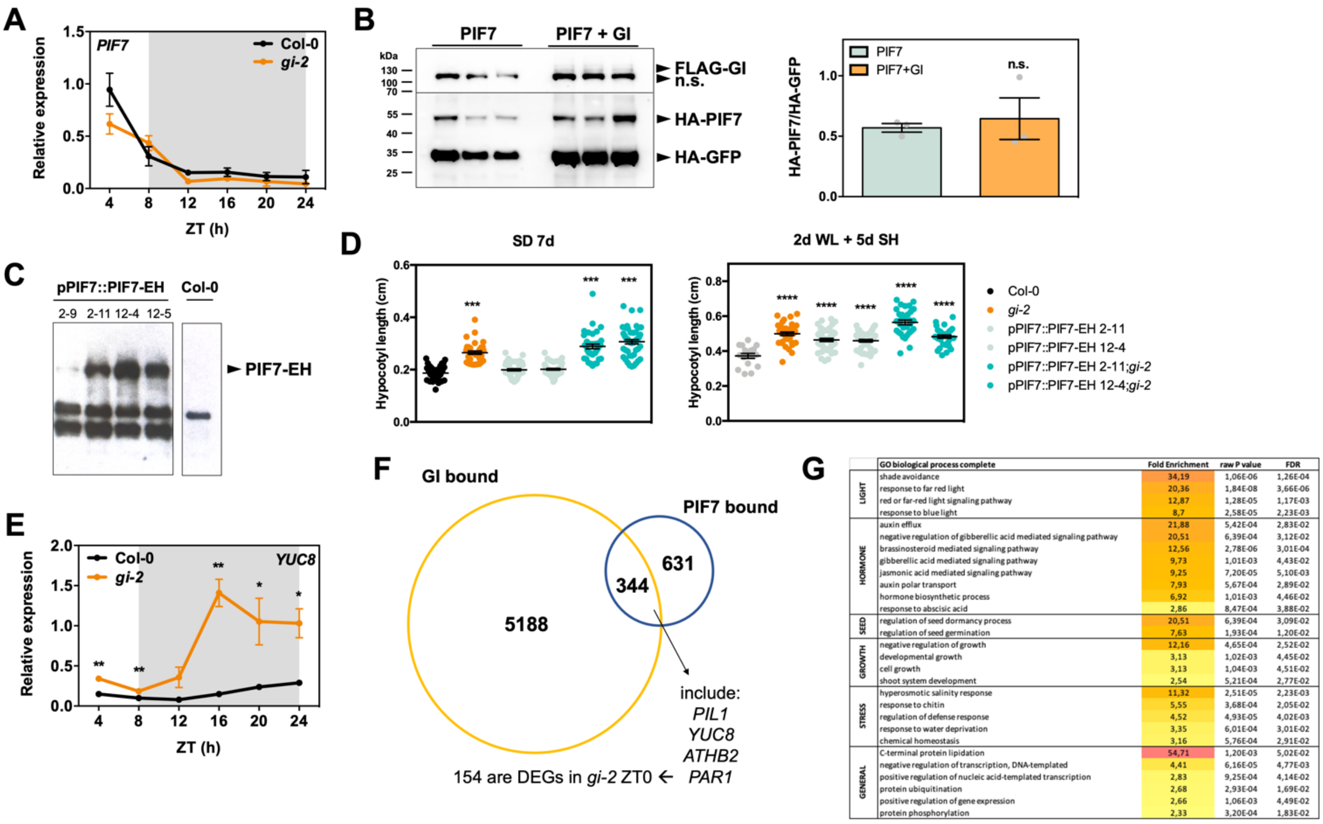


**Figure S3.**

(*A*) Relative expression of *PIF7* in wildtype (Col-0) and *gi-2* seedlings grown for 10 days in SDs (mean ± SEM of 3 biological replicates). White and gray shadings represent day and night, respectively. (*B*) HA-PIF7 protein accumulation in *N. benthamiana* leaves in the presence or absence of FLAG-GI. Protein levels were normalized against HA-GFP levels (n.s. not specific). Western blot quantitation is shown on the right panel (values represent mean ± SEM (n=3); n.s. not significant Student's *t* test). (*C*) Western blot analysis of the expression of PIF7-ECFP-HA from an endogenous promoter fragment (pPIF7::PIF7-ECFP-HA) in two independent transgenic lines (#2 and 12). Protein levels were determined with anti-HA antibody. (*D*) Physiological characterization of the pPIF7::PIF7-ECFP-HA transgenic lines. Hypocotyl length measurements from the indicated genotypes were grown for 7 days under SD conditions (left panel) or under continuous white light for 2 days and then transferred to constant shade light for 5 days (right panel). Mean ± SEM, n=17-54; ****p<0.0001, ***p<0.001, **p<0.01, *p<0.05 Tukey's multiple comparison test. (*E*) Relative expression of *YUC8* in wildtype (Col-0) and *gi-2* seedlings grown for 10 days in SDs (mean ± SEM of 3 biological replicates). White and gray shadings represent day and night, respectively. **p<0.01, *p<0.05 Student's *t* test. (*F*) Overlap between GI (19) and PIF7 (32) bound genes (hypergeometric test p value < 1.230e-23). (*G*) Heat map showing the GO category enrichment scores of genomic targets shared by GI and PIF7.

A

| | GO biological process complete | Fold Enrichment | raw P value | FDR |
|---|---|---|---|---|
| GROWTH | cell-cell junction assembly | 40,7 | 1.41E-04 | 1.12E-02 |
| | suberin biosynthetic process | 18,5 | 1.66E-05 | 1.74E-03 |
| | plant-type cell wall loosening | 13,2 | 1.28E-05 | 1.45E-03 |
| | xyloglucan metabolic process | 9,66 | 1.54E-05 | 1.65E-03 |
| | unidimensional cell growth | 3,83 | 4.43E-06 | 6.07E-04 |
| | cell wall biogenesis | 3,63 | 1.39E-06 | 2.28E-04 |
| OTHER DEV | stamen filament development | 29,6 | 2.62E-05 | 2.45E-03 |
| | gravitropism | 5,87 | 2.83E-04 | 1.94E-02 |
| | root development | 2,25 | 2.25E-05 | 2.25E-03 |
| | plant organ morphogenesis | 2,15 | 6.13E-04 | 3.57E-02 |
| HORMONES | auxin export across the plasma membrane | 18,09 | 1.31E-04 | 1.07E-02 |
| | auxin homeostasis | 10,17 | 2.10E-04 | 1.49E-02 |
| | auxin polar transport | 9,3 | 4.86E-06 | 6.50E-04 |
| | response to auxin | 8,42 | 3.99E-23 | 1.17E-19 |
| | response to brassinosteroid | 6,71 | 4.35E-05 | 3.77E-03 |
| | response to jasmonic acid | 2,62 | 1.27E-04 | 1.05E-02 |
| LIGHT | response to far red light | 12,66 | 3.05E-06 | 4.61E-04 |
| | phototropism | 11,63 | 5.86E-04 | 3.49E-02 |
| | response to red light | 10,48 | 1.42E-09 | 4.92E-07 |
| | photomorphogenesis | 6,63 | 1.41E-04 | 1.11E-02 |
| | response to light intensity | 4,38 | 3.94E-08 | 1.16E-05 |
| | cellular response to light stimulus | 4,28 | 7.83E-04 | 4.24E-02 |
| METABOLISM | olefinic compound biosynthetic process | 10,44 | 1.88E-04 | 1.40E-02 |
| | organic hydroxy compound biosynthetic process | 3,72 | 5.23E-04 | 3.25E-02 |
| | lipid biosynthetic process | 2,56 | 3.81E-05 | 3.46E-03 |
| | carboxylic acid metabolic process | 1,81 | 8.73E-04 | 4.68E-02 |
| | RNA metabolic process | 0,24 | 3.33E-04 | 2.23E-02 |
| | cellular nitrogen compound biosynthetic process | 0,19 | 2.06E-04 | 1.48E-02 |
| | proteolysis involved in cellular protein catabolic process | <0,01 | 7.05E-04 | 3.92E-02 |
| STRESS | response to oxidative stress | 3,32 | 3.36E-07 | 7.06E-05 |
| | response to cold | 3,21 | 2.38E-05 | 2.33E-03 |
| | response to hypoxia | 3,01 | 9.26E-04 | 4.91E-02 |
| | response to wounding | 2,72 | 2.88E-06 | 4.58E-04 |
| OTHER | carboxylic acid transmembrane transport | 8,14 | 5.41E-04 | 3.32E-02 |
| | gene expression | 0,19 | 1.53E-05 | 1.67E-03 |

B

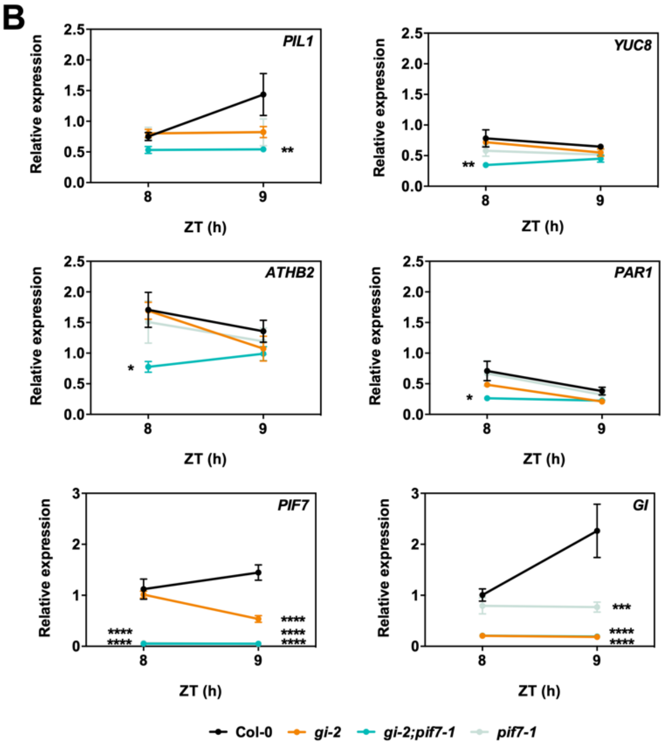


**Figure S4.**

(*A*) Heat map showing the GO category enrichment scores of the overlap between PIF7-dependent genes (32) and genes differentially expressed in *gi-2*. (*B*) White light controls to the experiment shown in Figure 4C. Relative expression of *PIL1*, *YUC8*, *ATHB2*, *PAR1*, *PIF7*, and *GI* in the indicated backgrounds kept in light and harvested at ZT 9. Seedlings were grown for 7 days under 10 h light/ 14 h dark photocycles. Mean ± SEM of 3 biological replicates. ****p<0.0001, ***p<0.001, **p<0.01, *p<0.05 Tukey's multiple comparison test.

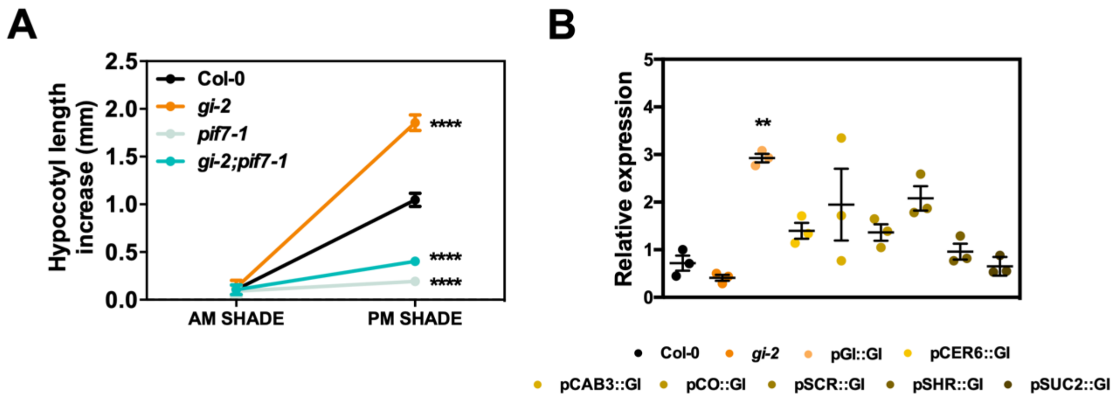


**Figure S5.**

(*A*) Hypocotyl length increase (measured as the difference between shade treated and white light kept seedlings) of seedlings grown for 3 days under 10 h light/ 14 h dark conditions and either exposed every day to 2 h shade light in the morning (AM SHADE, ZT 0 to 2) or in the afternoon (PM SHADE ZT 8 to 10) (mean ± SEM, n=10-18) (****$p<0.0001$ Tukey's multiple comparison test). (B) Relative expression of *GI* in the lines indicated grown for 7 days in SDs and harvested at ZT 8 (mean ± SEM of 3 biological replicates).
